# Transgenic tools for targeted chromosome rearrangements allow construction of balancer chromosomes in non-*melanogaster Drosophila* species

**DOI:** 10.1101/2021.11.18.469106

**Authors:** David L. Stern

**Author notes:** Corresponding Author: David L. Stern, Janelia Research Campus, Howard Hughes Medical Institute, 19700 Helix Dr, Ashburn, VA 20147, Tel: 571+209+4237.

## Abstract

Perhaps the most valuable single set of resources for genetic studies of *Drosophila melanogaster* is the collection of multiply-inverted chromosomes commonly known as balancer chromosomes. Balancers prevent the recovery of recombination exchange products within genomic regions included in inversions and allow perpetual maintenance of deleterious alleles in living stocks and the execution of complex genetic crosses. Balancer chromosomes have been generated traditionally by exposing animals to ionizing radiation and screening for altered chromosome structure or for unusual marker segregation patterns. These approaches are tedious and unpredictable, and have failed to produce the desired products in some species. Here I describe transgenic tools that allow targeted chromosome rearrangements in *Drosophila* species. The key new resources are engineered reporter genes containing introns with yeast recombination sites and enhancers that drive fluorescent reporter genes in multiple body regions. These tools were used to generate a doubly-inverted chromosome 3R in *D. simulans* that serves as an effective balancer chromosome.

## INTRODUCTION

Balancer chromosomes are inverted chromosome segments that prevent inheritance of the products of recombination events and thus allow maintenance of deleterious mutations in stable stocks (Hentges and Justice 2004; Kaufman 2017). A single chromosomal inversion can serve as a balancer chromosome, especially for alleles located close to inversion breakpoints. But balancer chromosomes have traditionally been constructed through serial introduction of new inversions in the context of previous inversions, because overlapping and nested inversions prevent inheritance of products of double recombination events (Kaufman 2017). Multiply inverted chromosomes have been the basis of all widely used balancer chromosomes in *D. melanogaster*, giving rise to the prefixes of many widely used balancers (FM = First Multiple, SM = Second Multiple, TM = Third Multiple) and balancers have facilitated many *D. melanogaster* experiments.

In plants, *C. elegans*, and mouse and human cells, inversions have been generated using CRISPR-Cas9 mutagenesis to induce simultaneously two double-strand breaks, which occasionally resolve as an inversion (Li *et al*. 2015; Iwata *et al*. 2016, 2019; Zhang *et al*. 2017; Korablev *et al*. 2017; Dejima *et al*. 2018; Schmidt *et al*. 2019). Similar efficient approaches have not been reported in *Drosophila*, and I have failed to recover inversions in *D. melanogaster* using this kind of approach to generate inversions on the order of 10kb-1Mbp. Recently, Ng and Reed reported successful isolation of a single inversion event in *D. melanogaster* using transgenic sources of Cas9 and gRNAs (Ng and Reed 2019). The efficiency of this approach is not yet known. The success of this method depended on a clever positive selection scheme that allowed survival of only animals that carried an inversion. It is not yet clear whether this method can be modified to incorporate markers or selection to allow identification of inversion events in a more general setting.

Inversions have traditionally been generated in *Drosophila* species by exposing flies to ionizing radiation that generates chromosome breaks, which are then incorrectly repaired. Balancer chromosomes would be valuable for diverse *Drosophila* species, but perhaps most useful for *D. simulans*, a sister species to *D. melanogaster*. Comparisons between *D. melanogaster* and *D. simulans* have provided fundamental insights into genetic evolution ever since *D. simulans* was discovered by Alfred Sturtevant in 1919 (Barbash 2010). However, while irradiation is effective to generate chromosome inversions in multiple species, for reasons that are not clear, it has proven difficult to induce inversions by mutagenesis in *D. simulans*. Several studies have found that irradiation and chemical mutagenesis induce rearrangements at a lower rate in *D. simulans* than in *D. melanogaster* (Lemke *et al*. 1978; Inoue 1988), although Woodruff and Ashburner (Woodruff and Ashburner 1975) observed irradiation-induced chromosome breaks at equal frequencies in each species, indicating that *D. simulans* retains mechanisms required for chromosome repair. In addition, inversion polymorphisms are nearly absent in *D. simulans* populations but common in *D. melanogaster* populations (Aulard *et al*. 2004). I am aware of several previous unpublished efforts to generate inversions in *D. simulans* using irradiation, but apparently none yielded useful balancer chromosomes.

One inversion stock is available in *D. simulans* with chromosome breaks at cytological locations 81F1 (approximately 0.91 Mbp on 3R) and 89E (9.75 Mbp), in the proximal region of chromosome 3R (Coyne and Sniegowski 1994). The distal break apparently disrupts *Ultrabithorax* (*Ubx*), since one breakpoint is at 89E, the cytological location of *Ubx*, and the inversion carries an *Ubx* allele. This inversion was probably generated by E. H. Grell (Coyne and Sniegowski 1994), perhaps using irradiation, though no details are available. This inversion can be stably maintained over a dominant homozygous lethal marker, *Delta*, located at approximately 7.15 Mbp, suggesting that it suppresses recombination at least between approximately 7 Mbp and 9.75 Mbp. However, this inversion does not serve as an effective balancer for several markers on 3R (Jones and Orr 1998), although the molecular locations of some of these markers are not known.

Here I describe new reagents developed to simplify the construction of overlapping inversions. The key innovations are the development of reporter genes with engineered introns containing yeast recombination sites and the expansion of the palette of enhancers used to drive fluorescent reporter genes in multiple body regions of the adult fly. The use of these reagents is illustrated by the construction of a *D. simulans* doubly-inverted 3R. This doubly-inverted chromosome arm prevented recombination on 3R over fifteen generations and thus will serve as an effective balancer chromosome for *D. simulans*. These transgenic reagents should work in multiple *Drosophila* species, thus democratizing the construction of balancer chromosomes, and other modified chromosomes, for non-*melanogaster Drosophila* species.

## METHODS

### Design

An autosomal balancer chromosome should contain multiple overlapping inversions, a dominant marker, and a recessive lethal allele (Kaufman 2017). It is straightforward to introduce recessive lethal alleles onto existing chromosomes with chemical mutagenesis or CRISPR mutagenesis, so I designed reagents that would accomplish the first two goals, a chromosomal region containing overlapping inversions and a dominant marker. I exploited a collection of existing mapped *pBac* transgenic *D. simulans* strains that each carry a dominant fluorescent marker (*3P3*-*EYFP*) adjacent to a site-specific *attP* recombination site (Stern *et al*. 2017). I integrated new reagents into these *attP* sites that would be modified during inversion events but leave the *3P3*-*EYFP* markers unaltered, ultimately generating inverted chromosomes carrying *3P3*-*EYFP* (Figure 1). I thus required enough markers to follow all of the transgenes in a single experiment. Generating a double inversion requires, at a minimum, six markers; one dominant marker that remains in the balancer chromosome; four markers to identify the sites targeted for recombination; and one marker for the recombinase. I performed these initial experiments in *D. simulans* flies carrying *w*- and used recombinase transgenes carrying mini-*white*. Since the YFP/GFP channel is occupied by the *attP* landing sites, only two other channels (blue and red) could be used to visualize fluorescent molecules easily in fly eyes on a binocular microscope. I therefore first searched for additional enhancers that would drive high levels of fluorescence in other body regions.

**Figure 1.**
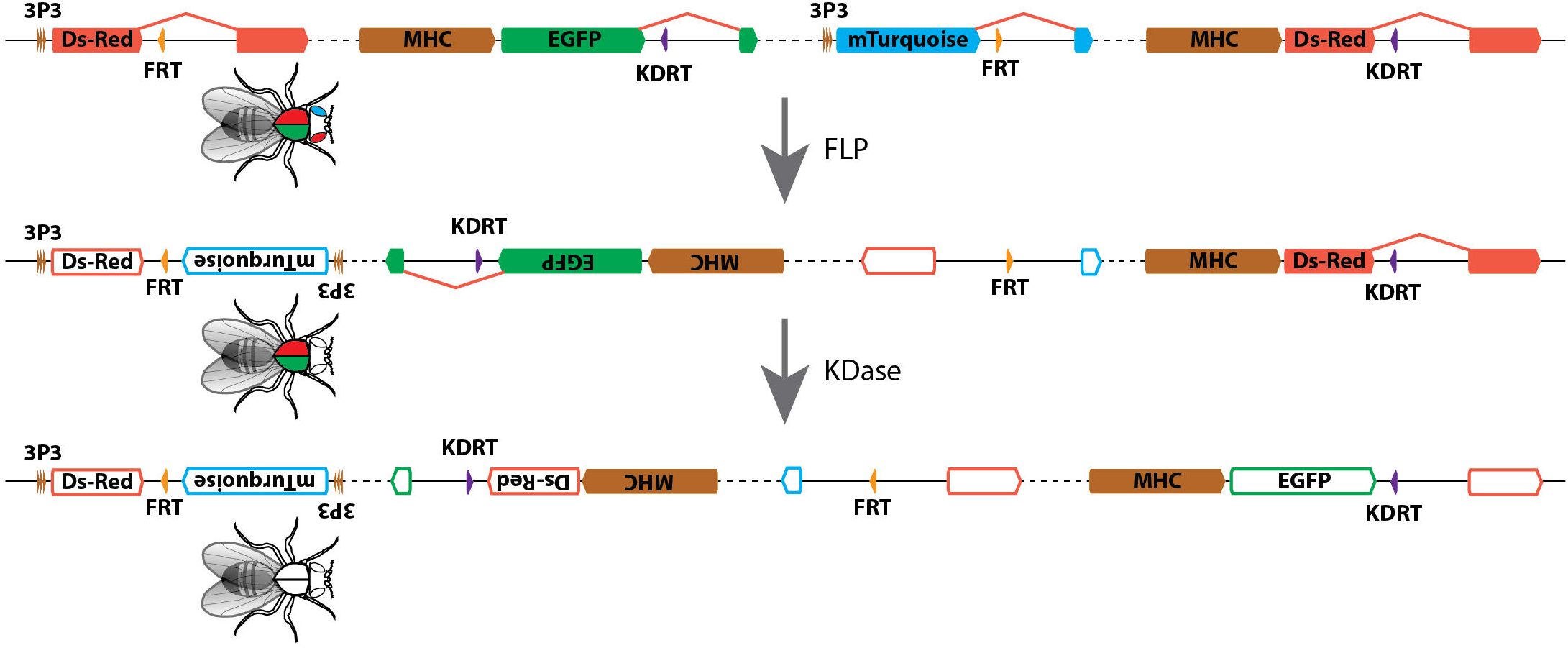
Illustration of the crossing scheme to generate targeted overlapping inversions. All four transgenes, each integrated into a separate landing site, are recombined onto a single chromosome arm. Dashed line represents long distances on chromosome. Filled color boxes represent genes encoding fluorescent proteins (red = DsRed, green = EGFP, blue = mTurquoise), enhancer sequences, or yeast recombination sites (orange = FRT, purple = KDRT). The yeast recombination sites are embedded within introns that have been engineered into reporter genes. Exposure of this chromosome to Flippase causes recombination between the FRT sites, generating an inversion that inactivates both the 3P3-DsRed and 3P3-mTurquoise genes. Inactivated genes are represented by geometric shapes with colored outlines and white fills. The first inversion event reverses the orientation of the MHC-EGFP∷KDRT reporter gene, resulting in two KDRT sites in inverted orientation on the chromosome. Exposure of this chromosome to KD recombinase can drive recombination between the KDRT sites, resulting in the second inversion and inactivation of the MHC-EGFP and MHC-DsRed genes. The 3P3-EYFP reporter genes in the landing sites remain after both inversion events and allow tracking of the doubly-inverted chromosome arm.

### New reporter constructs

Reporter constructs were generated to test the ability of synthetic combinations of characterized transcription factor binding sites for Tinman (Kimbrell *et al*. 2002) and Mef2 (Andres *et al*. 1995) to drive a fluorescent reporter in specific anatomical domains. gBlocks (IDT) were synthesized carrying 3 or 9 binding sites of each transcription factor binding site and homology arms complementary to the plasmid sequences, allowing Gibson cloning (Gibson *et al*. 2009) of these engineered enhancers upstream of eGFP reporter genes in *piggyBac* plasmids carrying an *attB* site-specific recombination sequence, allowing integration either through *piggyBac* transposition or *attP*-*attB* site-specific integration (Table S1). A second set of Tinman and Mef2 transcription factor binding site constructs were generated with GAGA binding sites (Wilkins and Lis 1998; Leibovitch *et al*. 2002) to attempt to boost expression levels (Table S1).

Reporter constructs were generated to test 145 bp and 285 bp enhancers of the *Actin88F* gene and 453 bp and 678 bp enhancers of the *Myosin Heavy Chain* gene (Table S1). The *ie1* enhancer (Masumoto *et al*. 2012) was PCR amplified from a plasmid provided by Rory Coleman and Gibson cloned upstream of fluorescent reporter genes.

The full list of plasmids generated for this project are listed in Table S2. All plasmids were re-sequenced in their entirety and annotated full-plasmid sequences are provided as Supplementary Material. All plasmids are available from Addgene.org.

### Generation of transgenic strains

Plasmids were introduced into the *D. melanogaster attP2* site-specific recombination site or various *D. simulans attP* sites by standard injection protocols performed by Rainbow Transgenics. For *D. simulans* injections, we compared co-injection of *attB* plasmids with a “helper” plasmid carrying phiC31 integrase under the control of a heat-shock inducible promoter versus mRNA encoding phiC31 integrase. PhiC31 integrase mRNA was generated as follows. p(T7-phiC31-nos-3’UTR) plasmid DNA was linearized by digestion with EcoRI and electrophoresed in a 1% agarose gel in 40mM Tris, 20mM Acetic Acid, 1mM EDTA buffer. The gel extracted linearized DNA was cleaned with the Zymoclean Gel DNA recovery kit (Zymo) and used as template in mMESSAGE mMACHINE™ T7 Transcription Kit (Thermo Fisher). RNA was cleaned with a LiCl_2_ precipitation followed by cleanup with a MEGAclear Transcription Clean-Up kit (ThermoFisher). Embryos were injected with 200 ng/uL of plasmid DNA and 500 ng/uL of mRNA.

### Reporter gene expression patterns

Photomicrographs of expression of reporter constructs in living adults were generated with an Olympus MBX10 macro zoom microscope with microscope magnification and exposure conditions held constant across all photographs.

A subset of the reporter lines were selected to examine potential reporter gene expression in the central nervous system. Central nervous systems were dissected, antibody stained, and imaged by Project Technical Resources at Janelia Research Campus as described previously (Aso et al. 2014). In brief, brains and VNCs were dissected in Schneider’s insect medium and fixed in 2% paraformaldehyde (diluted in the same medium) at room temperature for 55 min. Tissues were washed in PBT (0.5% Triton X-100 in phosphate buffered saline) and blocked using 5% normal goat serum before incubation with antibodies. DsRed was detected by staining with rabbit anti-dsRed (Clontech 632496, 1:1000) followed by CyTM3-conjugated goat anti-rabbit (Jackson ImmunoResearch 111-165-144, 1:1000). The neuropil was labeled with mouse anti-BRP hybridoma supernatant (nc82, Developmental Studies Hybridoma Bank, Univ. Iowa, 1:30), followed by CyTM2-conjugated goat anti-mouse (Jackson ImmunoResearch 115-225-166, 1:1000). After staining and post-fixation in 4% paraformaldehyde, tissues were mounted on poly-L-lysine-coated cover slips, cleared, and embedded in DPX as described previously. Image Z-stacks were collected at 1 um intervals using an LSM710 confocal microscope (Zeiss, Germany) fitted with a Plan-Apochromat 20x/ 0.8 M27 objective.

### Generation of a fluorescent reporter genes with engineered introns carrying Gateway cloning cassettes

To make plasmids that would allow flexible cloning of multiple reagents into engineered introns within fluorescent reporter genes, the Gateway cloning cassette (*attR2*-*ccdB*-*CAT*-*attR1*) was introduced into an intron cloned from the *D. melanogaster white* gene. This *white* intron-Gateway cassette, including the native 5’ splice donor and 3’ splice acceptor sites, was cloned into suitable sites (in between AG and GT) in the coding sequences for *mTurquoise* (Goedhart *et al*. 2010), *EGFP* (Cormack *et al*. 1996), and *DsRed* (Matz *et al*. 1999) reporter genes. FRT and KDRT yeast recombination sites (Nern *et al*. 2011) in forward and reverse orientations were cloned into a pENTR vector and plasmids used for site-specific chromosomal inversion were generated by Gateway cloning (ThermoFisher Scientific) of yeast recombination sites into introns of fluorescent reporter genes.

### *Generation of doubly-inverted* D. simulans *chromosome 3R*

Four plasmids carrying different reporter genes and yeast recombination sites were integrated separately into existing *attP* landing sites in *D. simulans* (Stern *et al*. 2017): p{3P3-dsRed∷FRT, attB} into 367-simD-311.3 at 3R:1,235,923 bp; p{Mhc-F4-678-EGFP∷KDRT-RC, attB} into 1392-simD-925.4 at 3R:10,884,926; p{3P3-mTurquoise∷FRT-RC, attB} into 1029-simD-234.5 at 3R:17,243,204; and p{Mhc-F4-678-DsRed∷KDRT, attB} into 1042-simD-275.5 at 3R:27,843,913. All genome coordinates are relative to *D. simulans* genome assembly NCBI:GCA_016746395.1. All four transgenes were recombined onto a single chromosome arm. Flies carrying all four transgenes on a single chromosome were crossed to flies carrying a source of heat-shock inducible Flippase p(MUH-HES-FLP1∷PEST, w+, attB) integrated into 1173-simD-523.7 at X:13,484,041. Plastic vials containing developing larvae in their food media were placed in a 37°C bead bath (Lab Armor) for 1 hour, followed by 1 hour at room temperature, and a second 37°C heat shock for one hour. Male progeny were crossed to a *w*^−^ strain of *D. simulans* and offspring of this cross were screened for loss of 3P3-DsRed and 3P3-mTurquoise, but retention of MHC-EGFP and MHC-DsRed, which indicates inversion between the FRT sites. Flies carrying this singly-inverted chromosome were then crossed to flies carrying a source of heat-shock inducible KD recombinase p(MUH-HES-KD∷PEST, w+, attB) integrated into 2176-simD-299.6 at 2L:6,583,842. This *attP* landing site is derived from 1048-simD-299.6, but has had its 3P3-EYFP marker inactivated by CRISPR-Cas9 mutagenesis of the EYFP gene (Stern *et al*. 2017). Larvae from this cross were heat shocked as described above, and adults were crossed to a *w*^−^ strain of *D. simulans*. Offspring were screened for loss of MHC-EGFP and MHC-DsRed, but presence of 3P3-EYFP, which is present in all four landing sites with integrated recombination plasmids, but not in the landing site with the KD-recombinase integrated. These progeny carried presumptive double-inversions on chromosome 3R.

### Testing for recombination with balancer chromosome

To determine whether the doubly-inverted chromosome arm might serve as a useful balancer chromosome, flies carrying this chromosome, which could be recognized by the 3P3-EYFP markers, were backcrossed to a *w*^−^ strain of *D. mauritiana* for 15 generations. Ten independent backcrosses were performed and DNA was prepared from one individual of the 15^th^ generation from each backcross and subjected to multiplexed-shotgun genotyping to estimate chromosome ancestry (Andolfatto *et al*. 2011).

## RESULTS

### *An expanded palette of enhancers useful for fluorescent reporters in* Drosophila

Tracking multiple transgenes, especially in species with few existing genetic reagents, requires development of new markers. For multiple reasons, I favored use of fluorescent markers. Fluorescent markers can be used in many species (Stern *et al*. 2017), they can be used in flies with normal eye color (Horn and Wimmer 2000), and they are unlikely to disrupt behavior, whereas eye color and pigmentation markers commonly used in *D. melanogaster* disrupt behavior (Kyriacou *et al*. 1978; Zhang and Odenwald 1995; Suh and Jackson 2007; Anaka *et al*. 2008; Massey *et al*. 2019).

I first attempted to engineer synthetic enhancers similar to the 3P3 enhancer, which consists of three binding sites for the *Drosophila* Eyeless transcription factor, called P3 sites (Sheng *et al*. 1997). The 3P3 enhancer drives strong expression in the eyes and has become a popular enhancer to mark transgenes in many non-model species (Horn *et al*. 2000, 2002). As expected, the 3P3 enhancer drove strong expression in neurons of the optic lobe and in many other regions of the brain and ventral nervous system (Figure S1A-D), which may limit its utility for some purposes.

I first generated concatemers of transcription factor binding sites for the Tinman and Mef2 transcription factors, which I hypothesized might drive expression in the heart and the muscles, respectively (Andres *et al*. 1995; Kimbrell *et al*. 2002). The Tinman binding site constructs did not drive strong expression of fluorescent reporters that could be detected in adults (Figure 2A,B), even after addition of GAGA binding sites (Leibovitch *et al*. 2002) which were hypothesized to open chromatin to allow improved access from Tinman transcription factors (Figure 2C). The Mef2 binding site 3X and 9X concatemers drove weak expression in a subset of flight and leg muscles (Figure 2D,E). The 3XMef2 constructs also drove extensive expression in the nervous system (Figure S1E-H). These synthetic enhancers did not drive adult expression that was judged useful for most experiments.

**Figure 2.**
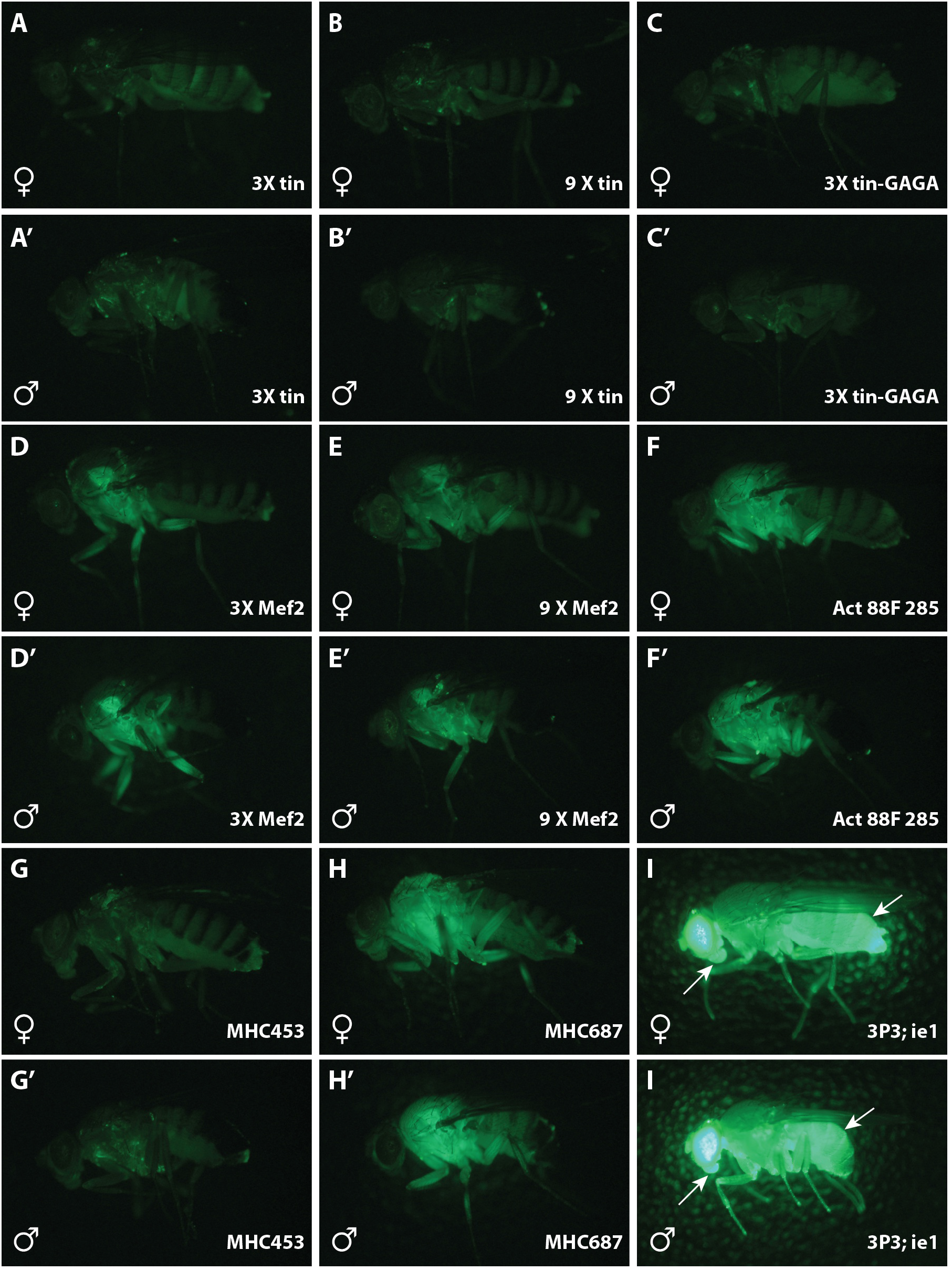
Adult expression of new reporter constructs. Lateral views of adult females (A-I) and males (A’-I’) expressing EGFP under control of tested reporter constructs. A-C – Reporter constructs driven by 3 (A) or 9 (B) multimerized Tinman transcription factor binding sites, or 3 Tinman sites combined with GAGA transcription factor binding sites did not result in observable fluorescence. D-E – Reporter constructs driven by 3 (A) or 9 (B) multimerized Mef2 transcription factor binding sites drove detectable, but relatively weak fluorescence in flight muscles. F – The *Act88F* 285 bp enhancer drove easily detectable expression in thorax and legs, and weak expression in eyes that is not visible in these images. G-H – The *MHC* 453 bp and 687 bp enhancers drove very weak and strong expression, respectively, in thoracic tissues. Of the enhancers tested, *MHC* 687 drove the strongest expression in thoracic tissues. I – Fly carrying *3P3*-*EYFP* and *ie*-*EGFP* displays strong expression in eyes (due to 3P3-EYFP) and strong expression in abdomen and mouthparts (arrows) due to ie-GFP. All *ie1* reporter genes were integrated into *3P3*-*EYFP* landing sites, so no images are available of the *ie1* reporter along.

A 145 bp and 285 bp fragment from the *D. melanogaster Actin 88F* gene were tested. No transgenic animals were recovered with the 145bp enhancer, suggesting that this enhancer did not support strong expression. The 285 bp enhancer drove strong expression in thoracic muscles (Figure 2F), but also drove some weak expression in eyes (which is not easily seen in Figure 2F). This enhancer drove no detectable expression in neurons of the brain or ventral nervous system in either sex (Figure S1I-L).

Enhancer fragments upstream of the *D. melanogaster myosin heavy chain* (*mhc*) gene have been shown to drive reporter gene expression in somatic muscles (Gajewski and Schulz 2010). A 453 bp *mhc* fragment drove weak expression that was barely detectable under a binocular microscope (Figure 2G). A 678 bp *mhc* fragment, however, drove strong expression in thoracic muscles in adults (Figure 2H) and was also visible in larvae. This enhancer drove stochastic expression in some neurons, especially a single mesothoracic motoneuron (Figure S1M-T).

I also tested the promoter of an immediate-early gene (*ie1*) isolated from a *Bombyx mori* nucleopolyhedrovirus, which had previously been shown to drive expression in *Drosophila* (Masumoto *et al*. 2012). This promoter drives strong expression in the gut and is visible in the abdomen and mouthparts of adult flies (Figure 2I).

The anatomical domains labeled by *3P3*, *mhc_678*, and *ie1* can be completely and rapidly distinguished in living flies (Figure 2 and S3), allowing, in principle, up to nine transgenes to be followed in a single fly using three genetically encoded fluorescent proteins visible in the blue, green, and red channels. These enhancers have been used to construct the reagents listed in Table S2 and a subset of these reagents were used to generate a targeted inversion, described below.

### *Generation of a multiply-inverted* D. simulans *chromosome 3R*

One way to track inversion events is to force recombination between the introns of two reporter genes located in *trans* (Egli *et al*. 2004). I therefore introduced yeast recombination sites into engineered introns within the coding sequences for *mTurquoise*, *EGFP*, and *DsRed* and integrated constructs driving different fluorescent reporters in different anatomical locations at four sites across chromosome 3R (Figure 1).

I found that almost all *attP* sites tested can support *phiC-31* induced site-specific integration if the source of integrase is provided as a mature mRNA transcript instead of as a heat-shock inducible “helper” plasmid (Figure S2). This finding allowed use of almost any *D. simulans attP* site in our collection for these experiments.

To drive separate inversions in two steps, I used different yeast recombination systems that are catalyzed by different recombinases (Nern *et al*. 2011). In this study I used the FRT/FLP and KDRT/KDase recombination systems. Inversion events were rare, but multiple independent events were isolated at each step after screening approximately 5,000-10,000 flies. Five independent doubly-inverted chromosomes were isolated and have the genotype In(3R){[367-1029][1392-1042], 3P3-EYFP+}. This doubly-inverted chromosome in *D. simulans* is called *j3RM1* (Janelia 3R Multiple 1).

### Confirmation of balancer efficiency

To determine the efficiency of *j3RM1* to balance the third chromosome, *D. simulans* flies carrying *j3RM1* were crossed to *D. mauritiana*, and virgin female offspring were backcrossed to *D. mauritiana* for 15 generations (Figure 3B). Crosses were made to *D. mauritiana* because *D. simulans* and *D. mauritiana* chromosomes are homosequential and display recombination throughout the genome (True *et al*. 1997). In addition, the species genomes are sufficiently divergent that even small genomic regions from each species can be identified unambiguously (Meiklejohn *et al*. 2018). Flies from ten independent backcrosses were subjected to light shotgun genome resequencing and ancestry of genomic regions was estimated using Multiplexed Shotgun Genotyping (Andolfatto *et al*. 2011). Flies from all ten crosses displayed heterozygosity for *D. simulans* and *D. mauritiana* DNA across the entire right arm of chromosome 3 (Figure 3B), indicating that *j3RM1* prevented recovery of recombination events on 3R for 15 generations.

**Figure 3.**
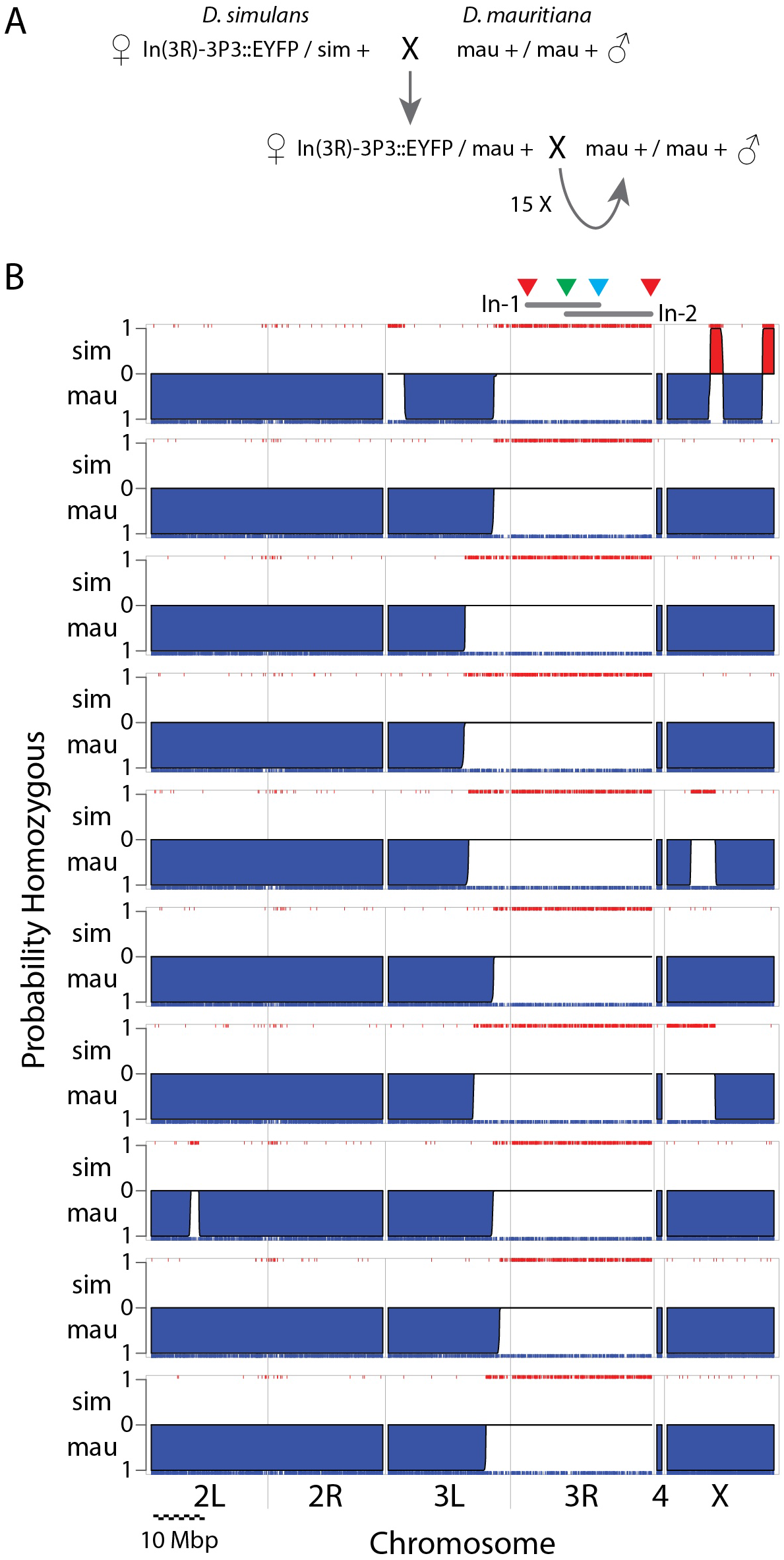
Testing for suppression of recombination by balancer j3RM1. A – *D. simulans* flies carrying j3RM1 were crossed to a *w*^−^ strain of *D. mauritiana*. Females of subsequent generations were backcrossed for 15 generations. B – Chromosomal ancestry estimates for single individuals from ten independent 15-generation backcross experiments. DNA from each individual was subjected to light shotgun sequencing and ancestry was estimated using the Hidden Markov Model of MSG (Andolfatto *et al*. 2011). The Y axis of each plot represents the probability that a chromosomal region is homozygous for either *D. simulans* (up, red regions) or *D. mauritiana* (down, blue regions). Individual DNA markers are shown as red and blue carpets along the top and bottom axes. Regions without support for homozygosity for either species are estimated to be heterozygous for the two species. Above the top plot, the approximate locations of the four markers (colored triangles) and two inverted segments (grey bars, labeled as In-1 and In-2) are indicated. In all individuals, the entire right arm of 3R is heterozygous, indicating the absence of recombination events within this region represented in progeny. In all individuals, the region of heterozygosity extends to the left of the third chromosome centromere, suggesting that the inversions on 3R suppress recombination at a considerable distance from the left-most breakpoint. The chromosome arms and scale bar in Mbp are indicated at bottom. For the scale bar, ten individual segments of 1 Mbp each are shown.

## DISCUSSION

The doubly-inverted *D. simulans* chromosome 3R, called *j3RM1*, prevented recombination over 15 generations in ten independent crosses (Figure 3B). This suggests that a single pair of overlapping inversions is sufficient to block inheritance of recombinant chromosomes to an extent that will make this balancer chromosome useful for many kinds of experiments. In our experience, *j3RM1* appears to have reduced viability relative to the un-inverted chromosome, but the chromosome is not homozygous lethal. However, it would be straightforward to add lethal alleles either through classical mutagenesis or CRISPR-Cas9 editing, and these manipulations are underway in our laboratory.

While the FRT and KDRT sites utilized here supported recombination between sites located approximately 16 and 20 Mbp apart, inversion events were rare. Although we did not quantify the inversion frequency precisely, we recovered approximately one inversion per several thousand flies. There may be several ways to increase the efficiency with which inversions are generated. For example, the heat-shock source of recombinases could be replaced with a stable germ-line source of recombinase. This might increase efficiency both by increasing the concentration of recombinase in cells and by exposing all germ cells to high levels of recombinase.

An alternative approach to increase efficiency might be to exploit homologous recombination to drive inversions. Egli et al. (2004) demonstrated that homologous recombination generated inversions in *D. melanogaster* at a frequency of 1-4%, independent of the distance between targeted sites. This is at least an order of magnitude more frequent than recombinase catalyzed inversions recovered in this study. Egli et al. targeted two double strand breaks using a rare-cutting restriction endonuclease that recognized a restriction site engineered into an intron of a *yellow* transgene. This general approach could be expanded by exploiting multiple rare-cutting restriction endonucleases (Taylor *et al*. 2012) or Cas9 gRNA sites. The target sites and long homology regions could be incorporated into the Gateway cloning sites of the plasmids described here.

### Alternative uses of these reagents

The fluorescent marker reagents described here simplified the generation of flies carrying many different transgenes, which we required to drive multiple inversions. One can envision many other potential uses of these markers. For example, many neurogenetic experiments performed in *D. melanogaster* require assembly of genotypes carrying many transgenes, most or all which are normally marked with mini-*white*. These complex crossing schemes are enabled, of course, by balancer chromosomes. However, the availability of up to nine dominant fluorescent reporters, using three colors to mark three body regions, may enable the construction of complex transgene genotypes in other species without requiring balancer chromosomes. The fact that the *act88F*, *MHC*, and *ie1* reporters drive limited expression in the nervous system (Figure S1) may make these more useful markers for neurogenetic studies than *3P3* reporters.

Most transgenic reagents used in *D. melanogaster* have employed visible markers that rescue eye color or cuticle pigmentation mutants. However, most or all of these mutations can influence development and behavior. For example, mini-*white* transgenes can influence male behavior (Zhang and Odenwald 1995; Anaka *et al*. 2008) and pigmentation mutations can alter behavior (Suh and Jackson 2007; Massey *et al*. 2019). Fluorescent markers may have fewer or weaker influences on development and behavior than transgenes carrying native *Drosophila* genes.

The reagents described here can be used to drive inversions, as illustrated, but could also be adapted to generate targeted deletions, as has proven useful in *D. melanogaster* (Ryder *et al*. 2004). The reagents can be adopted readily for other species carrying *attP* landing sites (Stern *et al*. 2017). Alternatively, the reagents could be introduced directly into targeted genomic locations using CRISPR-Cas9 homology directed repair to allow inversions and deletions of specific chromosomal regions. This might provide a means to simplify reversal of naturally-occurring inversions, making these regions accessible to further genetic studies (Schmidt *et al*. 2020).

## Supporting information

Plasmid Sequences

## ACKNOWLEDGEMENTS

I thank Rory Coleman for recommending use of the *ie1* promoter and for providing a plasmid containing the enhancer, Elizabeth Kim for making phiC31 mRNA and for assistance coordinating generation of transgenic flies, The Janelia Quantative Genomics team for re-sequencing all plasmids and generating the Illumina sequencing reads to assess genome ancestry, the Janelia Project Technical Resources team for staining and imaging fly nervous systems, and Richard Mann and Rory Coleman for discussion about the work.

---

**Figure S1.**
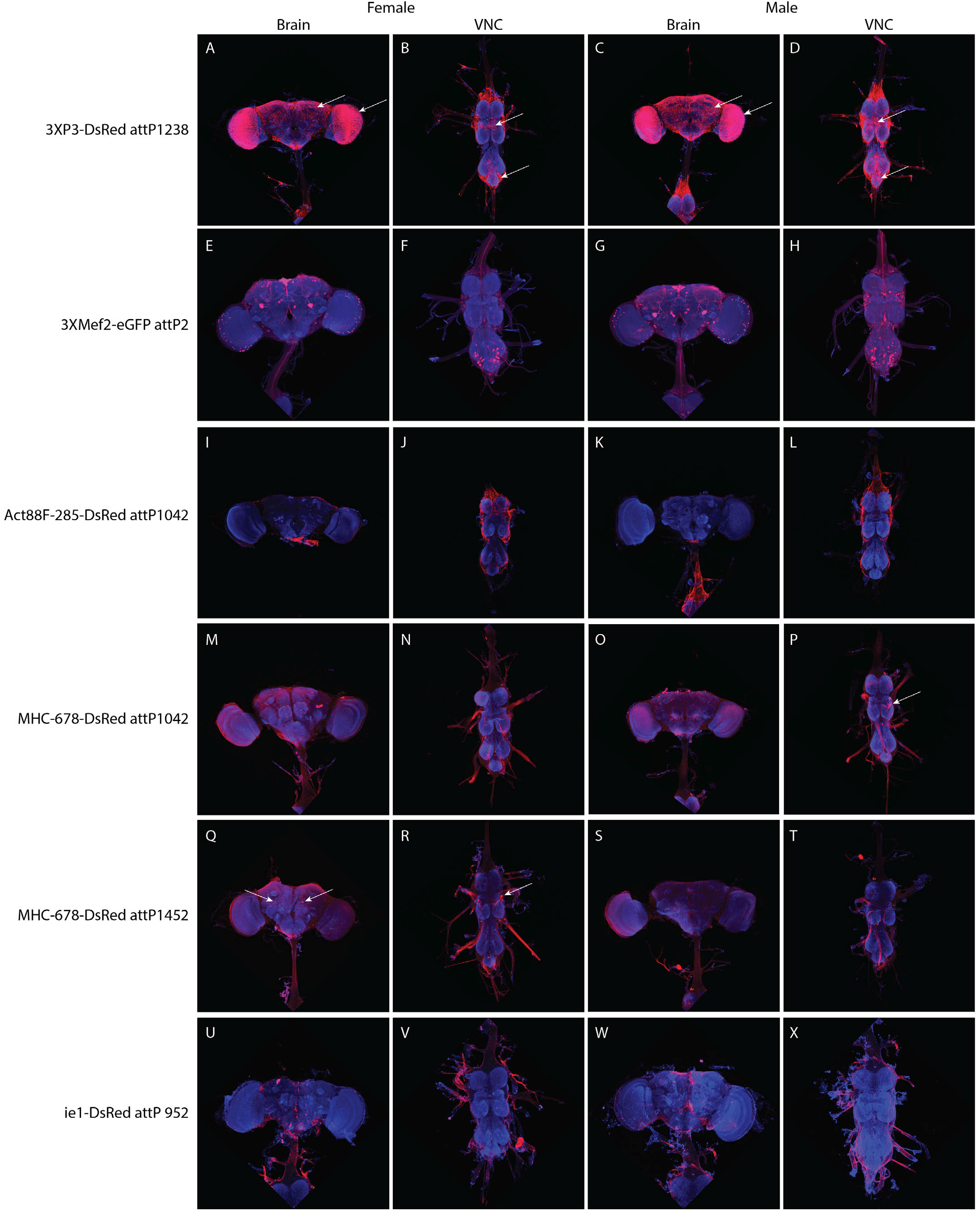
Expression patterns of fluorescent reporter genes in nervous system. Brains (columns 1 and 3) and ventral nerve cords (VNC; columns 2 and 4) from females (columns 1 and 2) and males (columns 3 and 4) stained for reporter genes (red) and neuropil (blue). A-D – Enhancer *3XP3* drives strong expression in optic lobe, as expected, but also in many other regions of the brain and VNC (white arrows point to regions of neuronal expression). E-H –Enhancer *3XMef2* drives expression in some neurons of the brain and ventral nerve cord and what appear to be glia. I-L – The *Act88F-285* enhancer did not drive detectable expression in brain or VNC cells. Red in these images is autofluorescence from non-neuronal tissue. M-T – The *MHC-687* enhancer in two different landing sites drove stochastic expression in a few neurons, most obviously a motor neuron of the mesothoracic segment. U-X – The *ie1* enhancer does not drive detectable expression in brain or VNC.

**Figure S2.**
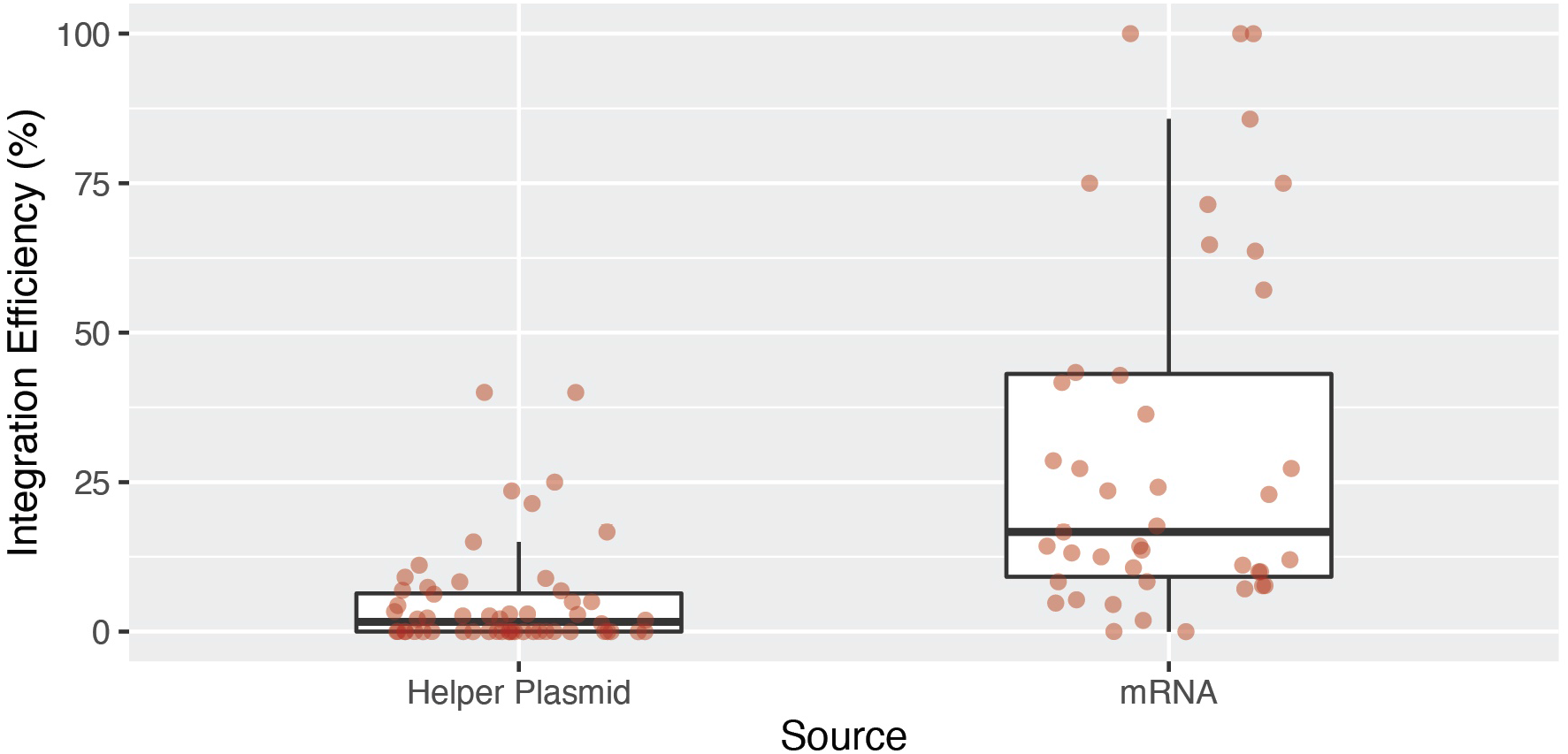
*In vitro* transcribed mRNA for phiC31 integrase provides increased integration efficiency compared with use of helper plasmid. Embryos carrying *attP* sites were injected with many different *attB* plasmids and co-injected with either pBS130 (56 injections), a “helper” plasmid carrying a heat-shock inducible *phiC31 integrase* gene, or *in vitro* transcribed *phiC31 integrase* mRNA (43 injections). Values indicate the proportion of fertile G0 animals that yielded offspring with integration events.

**Figure S3.**
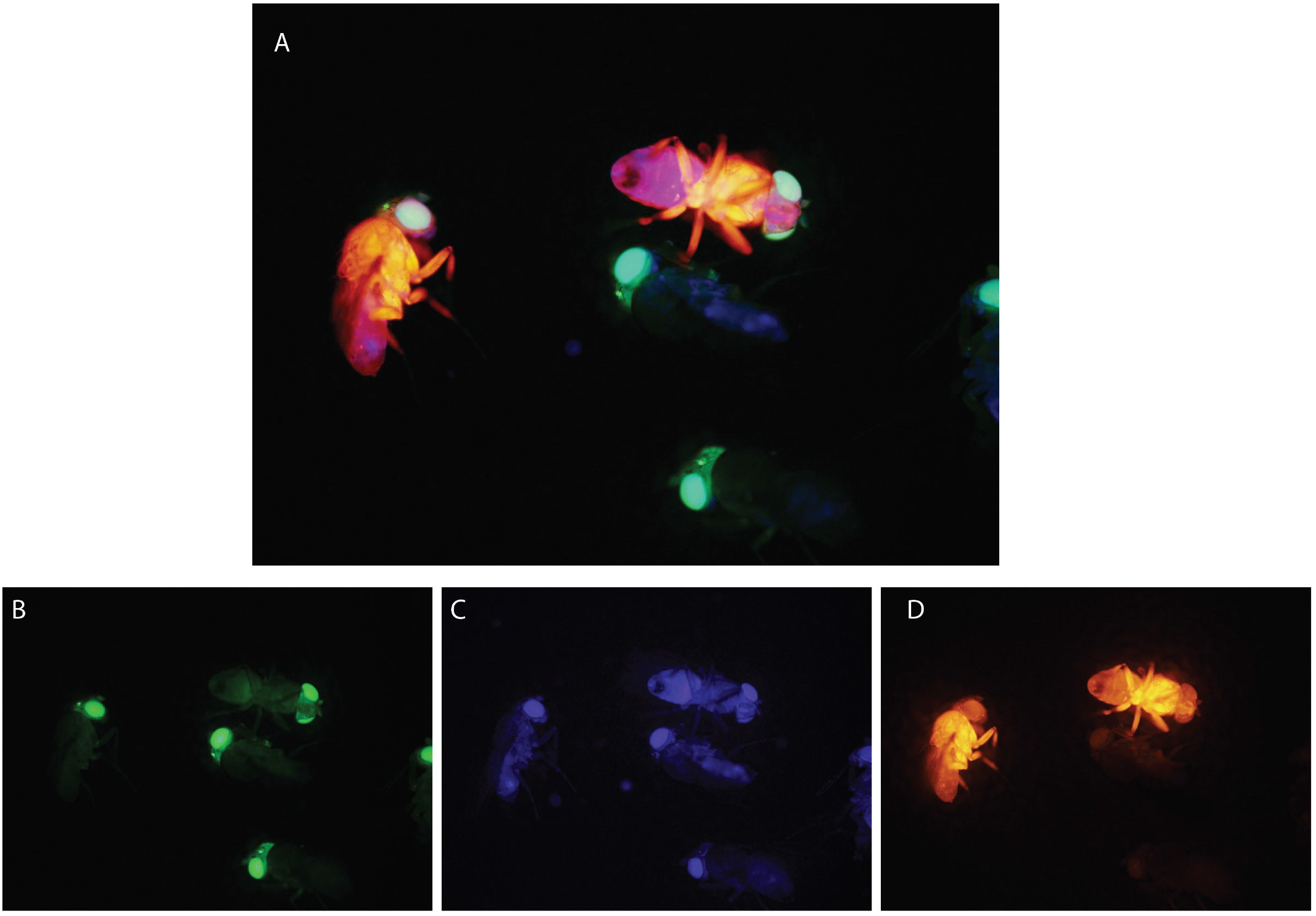
Example of flies segregating for three transgenes driving fluorescent proteins in two anatomical domains. (A) Flies expressing 3P3-EYFP and 3P3-mTurquoise and segregating for MHC-687-DsRed. (B-D) The three channels, green (B), blue (C), and red (D) illustrate that the colors and anatomical patterns can be clearly distinguished.

## SUPPLEMENTARY INFORMATION

Supplementary Material – All plasmids used in this study were re-sequenced in their entirety and annotated sequences are provided.

**Table S1.**
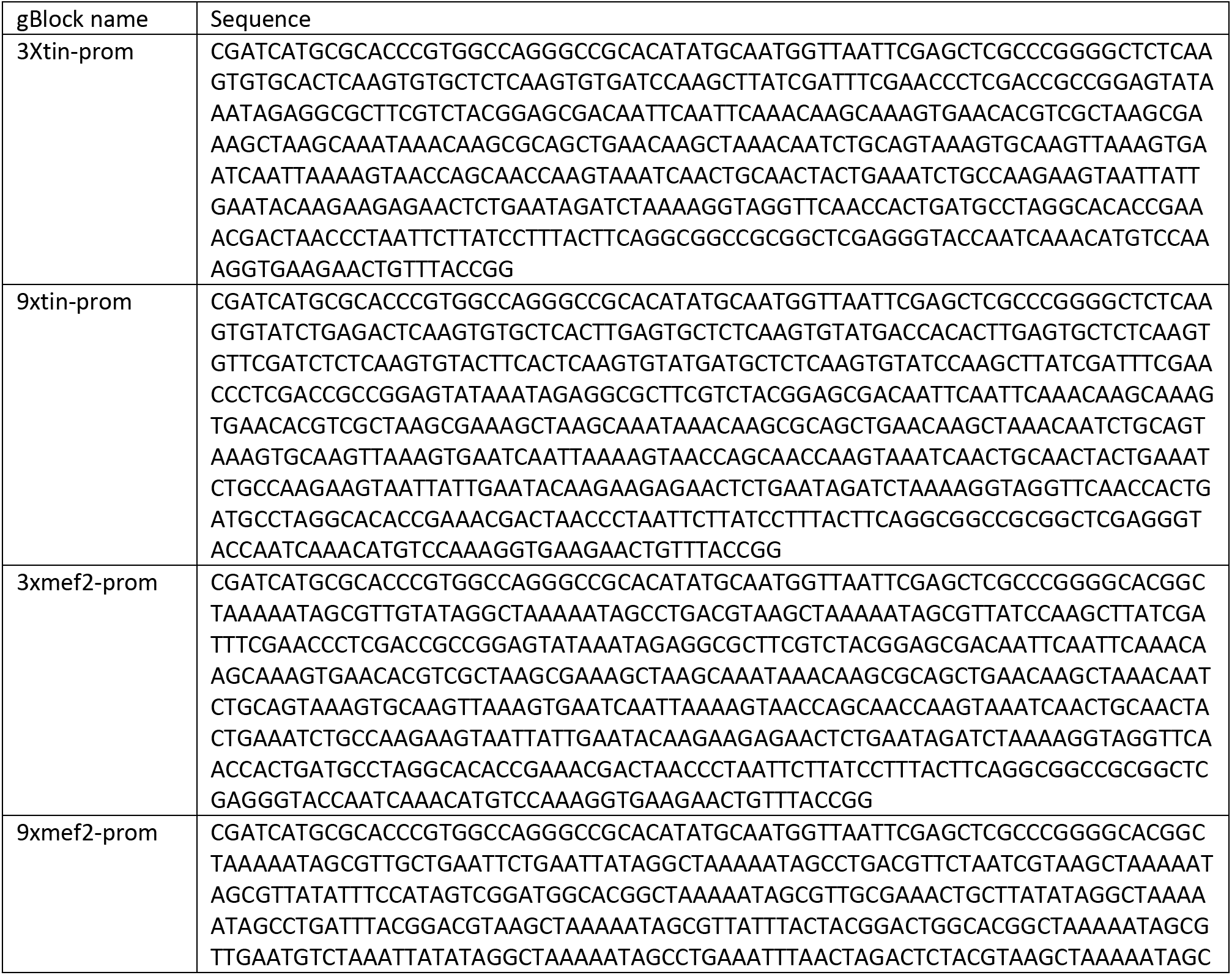
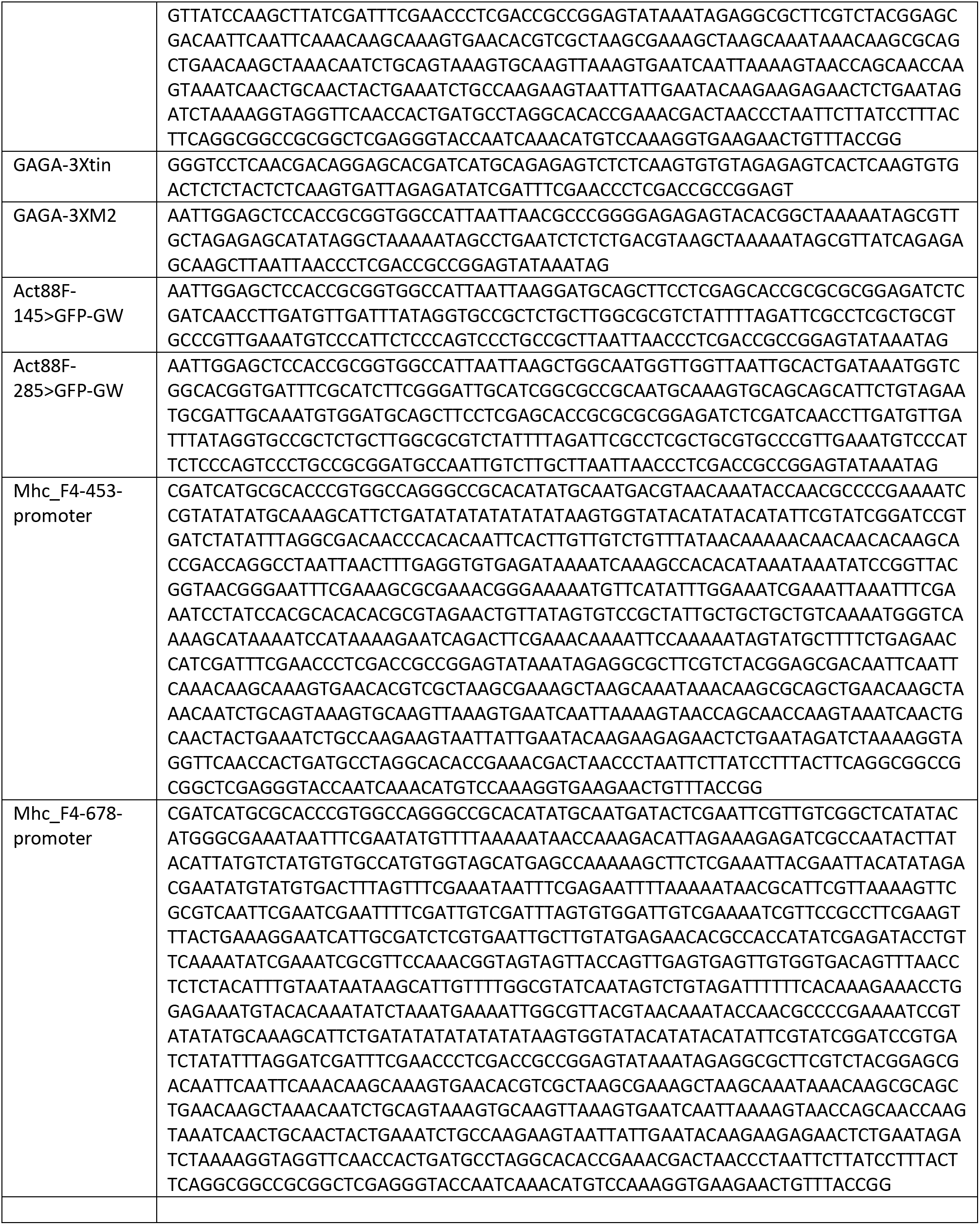
gBlocks used to generate new plasmids.

**Table S2.** Plasmids generated in this study. All plasmids are available from AddGene.

| Plasmid Name | AddGene Plasmid Number |
| --- | --- |
| pENTR{FRT} | TBD |
| pENTR{FRT-RC} | TBD |
| pENTR{KDRT} | TBD |
| pENTR{KDRT-RC} | TBD |
| p{3P3-eGFP::GW,attB} | TBD |
| p{3P3-dsRed::GW,attB} | TBD |
| p{3P3-mTurquoise::GW,attB} | TBD |
| p{Mhc-F4-678-EGFP::GW,attB} | TBD |
| p{Mhc-F4-678-DsRed::GW,attB} | TBD |
| p{ie1-eGFP::GW,attB} | TBD |
| p{ie1-DsRed::GW,attB} | TBD |
| p{ie1-eGFP::FRT,attB} | TBD |
| p{ie1-eGFP::FRT-RC,attB} | TBD |
| p{3P3-dsRed::FRT,attB} | TBD |
| p{3P3-dsRed::FRT-RC,attB} | TBD |
| p{3P3-dsRed::KD,attB} | TBD |
| p{3P3-dsRed::KD-RC,attB} | TBD |
| p{3P3-mTurquoise::FRT,attB} | TBD |
| p{3P3-mTurquoise::FRT-RC,attB} | TBD |
| p{3P3-mTurquoise::KD,attB} | TBD |
| p{Mhc-F4-678-eGFP::KD,attB} | TBD |
| p{Mhc-F4-678-GFP::KD-RC,attB} | TBD |
| p{Mhc-F4-678-eGFP::FRT,attB} | TBD |
| p{Mhc-F4-678-DsRed::FRT,attB} | TBD |
| p{Mhc-F4-678-DsRed::FRT-RC,attB} | TBD |
| p{Mhc-F4-678-DsRed::KD,attB} | TBD |
| p{Mhc-F4-678-DsRed::KD-RC,attB} | TBD |
| p{ie1-DsRed::FRT,attB} | TBD |
| p{ie1-DsRed::FRT-RC,attB} | TBD |
| T7-phiC31-nos3'UTR | TBD |
| pBac{3Xtin-eGFP,w+,attB} | TBD |
| pBac{9Xtin-eGFP,w+,attB} | TBD |
| pBac{3Xtin-GAGA-eGFP,w+,attB} | TBD |
| pBac{3Xmef2-eGFP,attB,w+} | TBD |
| pBac{9Xmef2-eGFP,attB,w+} | TBD |
| pBac{Act88F-eGFP,attB,w+} | TBD |
| pBac{Mhc-453-GFP,attB,w+} | TBD |

